# Effect of Light Conditions on *In Vitro* Adventitious Organogenesis of Cucumber Cultivars

**DOI:** 10.1101/2021.12.13.472500

**Authors:** Jorge Fonseca Miguel

## Abstract

The response on callus and shoot formation under different light incubation conditions was evaluated in cucumber (*Cucumis sativus* L.). Four-day-old cotyledon explants from the inbred line ‘Wisconsin 2843’ and the commercial cultivars ‘Marketer’ and ‘Negrito’ were employed. A four-week culture was conducted on MS-derived shoot induction medium containing 0.5 mg L-1 IAA and 2.5 mg L-1 BAP, under an 8-h dark/ 16-h light regime, or by a one- or two-week dark pre-incubation followed by the same photoperiod. Significant differences were obtained for the regeneration of shoots in all cultivars. The response in both frequency and number of shoots under continuous photoperiod was at least 3-6 fold higher than with dark pre-incubation. The highest genotypes response was obtained by ‘Negrito’ and ‘Marketer’ with identical values. All explants formed callus, and in two of the three cultivars, the response on callus extension was not significantly affected by incubation conditions. The results clearly show that shoot induction under continuous photoperiod regime was beneficial for adventitious shoot regeneration in cucumber.

## 1. Introduction

Cucumber (*Cucumis sativus* L.) belongs to the economically important family Cucurbitaceae along with melon, watermelon and squash. World production of cucumber, including gherkins, ranked in 2019 third among vegetable crops with 87,805,086 tonnes, and a harvested area of 2,231,402 hectares (FAOSTAT, http://www.fao.org/faostat/en/#data/QC). Crop improvement of cucumber for traits that confer resistance/tolerance to major biotic/abiotic stresses is difficult through conventional breeding due to its narrow genetic base, low genetic variability, and various crossing barriers with related species (Den Nijs and Custers 1990; Plader et al. 2007; Miguel 2021). Besides, conventional approaches are labor-intensive, time-consuming, and costly. Genetic engineering and plant transformation techniques have the potential to overcome these constraints (Miguel 2021). Despite the number of studies on genetic transformation of cucumber, its efficiency is still far from ideal (Wang et al. 2015; Miguel 2017). The main limitations in obtaining transgenic cucumber plants are the low morphogenetic response, inadequate selection methods, and the high rate of non-transgenic “escape” plants (Miguel 2017). By improving the efficiency of regeneration systems, we are addressing a part of the problem.

Regeneration via organogenesis and somatic embryogenesis has been reported in cucumber. Commonly used explants are cotyledons (Selvaraj et al. 2007; Grozeva and Velkov 2014; Miguel 2021), hypocotyls (Selvaraj et al. 2006; Grozeva and Velkov 2014), and leaves (Burza and Malepszy 1995a; Seo et al. 2000). Regeneration from protoplasts (Punja et al. 1990b; Burza and Malepszy 1995b) and suspension cultures (Raharjo and Punja 1994; Kreuger et al. 1996) has also been described. However, regeneration in this species is still not optimal (Wang et al. 2015), and is highly genotype-dependent (Wehner 1981; Punja et al. 1990a; Miguel 2021). An efficient and reproducible regeneration protocol is essential for successful genetic transformation of cucumber.

In *in vitro* plant regeneration, in addition to the culture factors that are usually considered, such as genotype, explant type, age of the donor plants, number and duration of subcultures, medium composition and growth regulators, the choice of appropriate incubation conditions, including light, temperature and humidity regimes, are essential to optimize regenerative responses.

Light, in particular, is a crucial environmental factor that, besides providing energy for photosynthesis, triggers and modulates complex developmental and regulatory processes (Tobin and Silverthorne 1985; Goins and Yorio 2000; Mawphlang and Kharshiing 2017). Plants can sense many parameters of environmental light, such as quality (spectral composition), light intensity, direction, and duration (including day length), and use this information to optimize growth and development during their whole life cycle (Chory 1997; Batschauer 1998; Heijde and Ulm 2012; Fiorucci and Fankhauser 2017). To sense and respond to environmental light conditions, plants are equipped with several classes of photoreceptors, other than photosynthetic pigments, including phytochromes, cryptochromes, phototropins, zeitlupe family members, and UVR-8, that can monitor specific ranges of the light spectrum (from UV-B to far-red), albeit with overlapping action spectra (Batschauer 1998; Kami et al. 2010; Galvão and Fankhauser 2015; Llorente et al. 2016). Plants have constantly to adapt to a varying light environment (Galvão and Fankhauser 2015). Despite their remarkable plasticity, light fluctuations can have a critical impact on plant competition and survival (Smith 2000; Paik and Huq 2019). The effect of light is most visible during seedling development. The patterns of seedling development under light (photomorphogenesis) differ from those under darkness (skotomorphogenesis or etiolation) regarding gene expression, differentiation, and organ morphology (Von Arnim and Deng 1996). Photomorphogenesis is characterized by short hypocotyls, open and expanded cotyledons, cell-type differentiation, chloroplast development, anthocyanin accumulation, and the expression of a large set of light-inducible genes encoded by the chloroplast and the nucleus. On the other hand, skotomorphogenesis is typically distinguished by long hypocotyls, closed and unexpanded cotyledons, closed apical hooks, and the development of etioplasts (Chory 1997; Jarillo and Cashmore 1998; Yang et al. 2005; Alabadí and Blázquez 2008; Qin et al. 2020). The interaction between environmental signals (light) and endogenous cues (gibberellin plant hormones, among others) determines the choice of one of the two processes (Alabadí and Blázquez 2008). The optimal lighting conditions depend on species, cultivars, plant growth stages, specific secondary metabolites, and other environmental parameters, such as nutrients, temperature, and CO_2_ levels (Dou and Niu 2020). The specific effects of light in a particular species can differ substantially between organs or cell types, even between nearby cells, as well as throughout development. (Von Arnim and Deng 1996). In *Arabidopsis* seedling, it was estimated that approximately 1/3 of the genes whose expression is regulated by light, where 3/5 are up-regulated and 2/5 are down-regulated (Ma et al. 2001), revealing, in particular, the crucial role of light and its complexity in the early stage of plant development. Light signaling pathways are interconnected with many other pathways to modulate plant physiology and development (Paik and Huq 2019).

This study aimed to determine the influence of different dark/light incubation regimes on *in vitro* adventitious organogenesis, using cotyledons as explants from one inbred line and two commercial cultivars of cucumber.

## 2. Materials and Methods

### 2.1. Plant material and regeneration

Cucumber (*Cucumis sativus*) seeds of the inbred line ‘Wisconsin 2843’ (courtesy of Dr. C.E. Peterson) and of the cultivars ‘Marketer’ and ‘Negrito’ (Semillas Fitó S.A.), were the starting material. Obtaining axenic explants and *in vitro* adventitious regeneration were based on the methodology previously described by Miguel (2021) in cucumber, with some modifications.

Seeds were surface-sterilized by immersion in a solution of 5% w/v sodium hypochlorite and 0.1 (v/v) 7X-O-matic (Flow Laboratories) for 30 min, and rinsed with sterile distilled water. They were then germinated on MS-derived medium without growth regulators. From 4-day-old axenic seedlings, cotyledons were excised and used as explant source by removing 1-2 mm behind their proximal and distal ends. All culture media were solidified with 0.8% (w/v) agar (Industrial, Pronadisa), and its pH adjusted to 5.7 before autoclaving. Plant material was incubated in a growth chamber at 26 ± 2 ºC under standard 16-h light/8-h dark photoperiod with cool-white fluorescent light at a photon fluence rate of 90 µmol m^−2^ s^−1^ (Grolux, Sylvania, fluorescent tubes); in dark pre-incubation, jars were wrapped with aluminum foil to prevent the passage of light. The experimental evaluations are based on observations with a naked eye.

Cotyledon explants were cultivated for 4 weeks on MS-derived shoot induction medium (SIM) containing 0.5 mg L-1 IAA and 2.5 mg L-1 BAP, under the standard photoperiod regime, or by a one- or two-week dark pre-incubation followed by the same photoperiod. Then, callus regeneration frequency (%) (CRF) and callus extension index (CEI) were determined, where CRF (mean±SE) is the frequency of explants with callus on the cutting zone and, CEI (mean±SE) correspond to arbitrary values (from 0 to 3) on the extension of callus on the cutting zone, where: 0 = absence of callus; 1= traces of callus; 2 = callus on less than half; 3 = callus on half or more; 4 = callus covering the full extension.

Adventitious buds and shoot primordia obtained were then cultured for 2 weeks on MS-derived shoot development and elongation medium (SDM) containing 0.2 mg L^-1^ KIN. Next, shoot regeneration frequency (%) (SRF) and shoot number index (SNI) were evaluated, where SRF (mean±SE) is the frequency of explants with shoots and, SNI (mean±SE) correspond to arbitrary values (from 0 to 3) on the number of shoots per explant, where: 0 = absence; 1 = one shoot; 2 = two shoots; 3 = three or more shoots. Individualized shoots were then rooted on hormone-free MS medium and were ready for acclimation in 3 to 4 weeks (data not shown).

### 2.2. Data analysis

The experiment was arranged in a 3 × 3 completely randomized factorial design. At least twelve replicate flasks of six explants each were used in each treatment. All statistics were carried out using R version 4.0.4 (R Core Team 2021). Non-linear regression analyses were used to compare treatment means. To perform logistic regression, COM-Poisson regression, and generalized Poisson regression, the R packages ‘stats’ (R Core Team 2021), ‘COMPoissonReg’ (Sellers et al. 2019), and ‘VGAM’ (Yee 2021), were used, respectively. Model fit was evaluated using Akaike Information Criterion (AIC; Akaike 1973) and Bayesian Information Criterion (BIC; Schwarz 1978). The level of statistical significance was set at P < 0.05.

## 3. Results

### 3.1. Frequency and extension of callus

Results are expressed as mean ± SEM (Table 1), with significance defined as P < 0.05. Callus was formed within the first two weeks of culture, starting at the cut ends of the primary explant. All explants formed callus (100% of frequency). Two of the three cultivars showed no significant differences on callus extension for the incubation conditions tested. In contrast, ‘Negrito’ cultivar showed differences between the 2-week dark pre-incubation treatment (2D/2F), the highest response (1.74±0.08), and the other incubation regimes, showing that including a longer pre-incubation favored callus extension. Other scores ranged from 1.44±0.06 to 1.58±0.06. The results lie between the arbitrary values on callus extension of 1-traces of callus, and 2-callus in less than half, at explant cut edges.

**Table 1.**
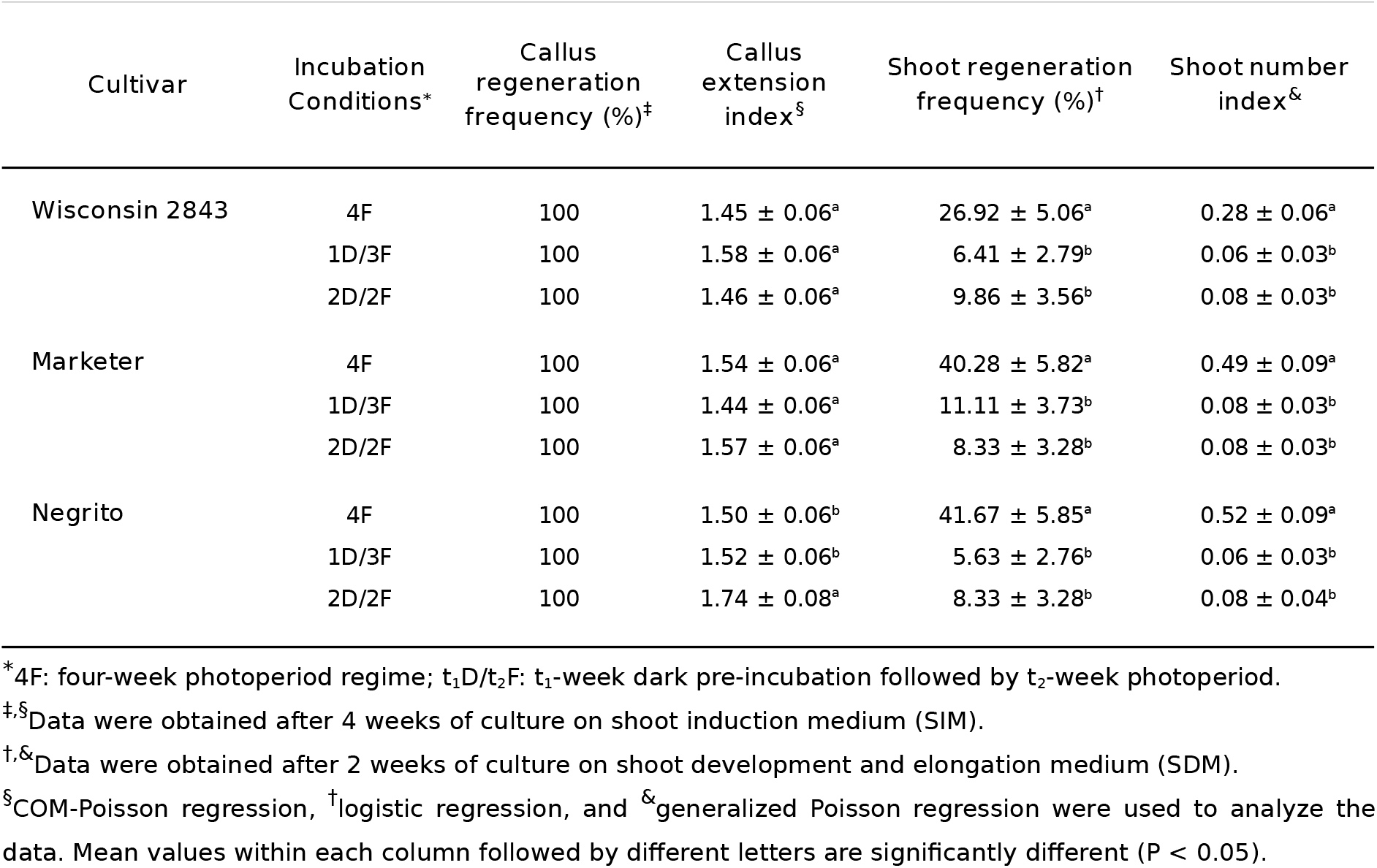
Effect of incubation conditions on *in vitro* callus and shoot regeneration from cotyledon explants of cucumber *(Cucumis sativus* L.) cultivars. Data are reported as mean ± standard error of the mean (SEM).

| Cultivar | Incubation Conditions* | Callus regeneration frequency (%)<sup>‡</sup> | Callus extension index<sup>§</sup> | Shoot regeneration frequency (%)<sup>†</sup> | Shoot number index<sup>&</sup> |
| --- | --- | --- | --- | --- | --- |
| Wisconsin 2843 | 4F | 100 | 1.45 ± 0.06<sup>a</sup> | 26.92 ± 5.06<sup>a</sup> | 0.28 ± 0.06<sup>a</sup> |
| | 1D/3F | 100 | 1.58 ± 0.06<sup>a</sup> | 6.41 ± 2.79<sup>b</sup> | 0.06 ± 0.03<sup>b</sup> |
| | 2D/2F | 100 | 1.46 ± 0.06<sup>a</sup> | 9.86 ± 3.56<sup>b</sup> | 0.08 ± 0.03<sup>b</sup> |
| Marketer | 4F | 100 | 1.54 ± 0.06<sup>a</sup> | 40.28 ± 5.82<sup>a</sup> | 0.49 ± 0.09<sup>a</sup> |
| | 1D/3F | 100 | 1.44 ± 0.06<sup>a</sup> | 11.11 ± 3.73<sup>b</sup> | 0.08 ± 0.03<sup>b</sup> |
| | 2D/2F | 100 | 1.57 ± 0.06<sup>a</sup> | 8.33 ± 3.28<sup>b</sup> | 0.08 ± 0.03<sup>b</sup> |
| Negrito | 4F | 100 | 1.50 ± 0.06<sup>b</sup> | 41.67 ± 5.85<sup>a</sup> | 0.52 ± 0.09<sup>a</sup> |
| | 1D/3F | 100 | 1.52 ± 0.06<sup>b</sup> | 5.63 ± 2.76<sup>b</sup> | 0.06 ± 0.03<sup>b</sup> |
| | 2D/2F | 100 | 1.74 ± 0.08<sup>a</sup> | 8.33 ± 3.28<sup>b</sup> | 0.08 ± 0.04<sup>b</sup> |

\* 4F: four-week photoperiod regime; t<sub>1</sub>D/t<sub>2</sub>F: t<sub>1</sub>-week dark pre-incubation followed by t<sub>2</sub>-week photoperiod.
<sup>‡,§</sup>Data were obtained after 4 weeks of culture on shoot induction medium (SIM).
<sup>†,&</sup>Data were obtained after 2 weeks of culture on shoot development and elongation medium (SDM).
<sup>§</sup>COM-Poisson regression, <sup>†</sup>logistic regression, and <sup>&</sup>generalized Poisson regression were used to analyze the data. Mean values within each column followed by different letters are significantly different (P < 0.05).

### 3.2. Frequency and number of shoots

Data are presented as mean ± SEM (Table 1), with significance set at P < 0.05. Shoots formed from callus within 2-4.5 weeks of culture, mainly at the proximal end of the explant. Those obtained in the treatments with pre-incubation in the dark were not etiolated. For both frequency (SRF) and shoot number index (SNI), cultivars followed the same pattern, with a 4-week photoperiod regime (4F) showing significant differences from the other treatments. Its response was at least 3-6 times greater than the dark pre-incubation treatments, which did not differ statistically from each other. With similar values to ‘Marketer’, ‘Negrito’ had the highest response, with a frequency of 41.67±5.85 and an index of 0.52±0.09. ‘Wisconsin’ has a 35% and a 46% lower frequency and index, respectively.

## 4. Discussion

The organogenic response of three cucumber cultivars was assessed under different lighting regimes: standard 16L:8D h photoperiod and two periods of dark pre-incubation.

Light is a critical environmental factor, that besides being the driving force behind photosynthesis, triggers and modulates complex developmental and regulatory processes (Tobin and Silverthorne 1985; Simpson and Herrera-Estrella 1990; Mawphlang and Kharshiing 2017). As an environmental cue, light regulates many aspects of plant biology. These include adaptive responses (e.g., phototropism, shade avoidance, and synthesis of photoprotective pigments), developmental transitions (e.g., germination, de-etiolation, flowering time, and senescence), and at the cellular level (e.g., chloroplasts movement, and stomatal opening) (Wang and Deng 2003; Meng et al. 2013; Takemiya et al. 2013; Galvão and Fankhauser 2015; Llorente et al. 2016; Vandenbussche et al. 2018). Light is also the main agent that mediates the entrainment of circadian rhythms (Arpaia et al. 1993).

In *in vitro* regeneration, explants are often incubated in a growth chamber under a 16L:8D h photoperiod. However, some researchers choose to perform a pre-incubation in the dark.

A dark pre-incubation has been reported in the micropropagation of different plant species. Regeneration via somatic embryogenesis and organogenesis with 2-3 weeks dark pre-treatment was described in two inbred lines and in an F1 hybrid of cucumber, using cotyledon, leaf, and petiole explants (Punja et al. 1990a). Regeneration via embryogenesis with 3-4 weeks pre-incubation was reported in cucumber cultivars from explants of petiole (Raharjo and Punja 1992), cotyledon and hypocotyl (Chee 1990). In other cucurbits, 2-3 weeks dark pre-treatment was described in the organogenesis of melon (*Cucumis melo* L.) and African horned cucumber (*C. metuliferus* E. Mey. ex Naudin) (Punja et al. 1990a), as well as in increasing somatic embryo production in melon and squash (*Cucurbita pepo* L.) (Kintzios et al. 1997). In species other than cucurbits, a pre-incubation in the dark has also been reported. A 2-3 weeks pre-treatment favored somatic embryo induction and development in pepper (*Capsicum annum* L.) (Kintzios et al. 1997). A 2-week dark pre-incubation was described in adventitious organogenesis of apple cultivars (Yepes and Aldwinckle 1994), and in increased frequency of somatic embryogenesis in purple coneflower (*Echinacea purpurea* (L.) Moench) (Zobayed and Saxena 2003). A 1-week pre-treatment enhanced shoot regeneration in chrysanthemum (*Chrysanthemum morifolium* Ramat) (Naing et al. 2015). In other reports, a dark pre-incubation had no positive effect on regeneration. Callus from cotyledon explants of cucumber failed to produce shoot buds in the dark (Selvaraj et al. 2007). A pre-incubation had a detrimental impact on gardenia (*Gardenia jasminoides* L.), and no somatic embryos were formed for less than 7 weeks in the dark (Kintzios et al. 1997). In rose (*Rosa hybrida* L.), a pre-treatment of 1 to 10 weeks in darkness failed to induce somatic embryogenesis (Kintzios et al. 1997).

In the present investigation, frequency and extension of callus were not influenced by light incubation conditions, except for cv. Negrito, where a 2-week dark pre-incubation enhanced callus extension. The findings on callus formation are in line with some reports. Punja et al. (1990a), when using different cucumber genotypes, growth regulators combinations, and explant types, the percentage of callus was not affected by pre-incubation in the dark. Likewise Gammoudi et al. (2017), when using pepper cotyledon explants, callus frequency was not influenced by a dark pre-treatment. The opposite was observed in hypocotyl explants. Also unlike the present study, in the organogenesis of bael (*Aegle marmelos* (L.) Corr.) using different explants, a dark incubation (1-7 days) favored abundant callus formation, with cotyledon explants and the 3-day treatment giving the best response (Arumugam et al. 2003). In anther culture of *Capsicum annuum* L., both growth regulators combinations and light regimes influenced the frequency and intensity of callus formation (Mythili and Thomas 1995).

The results on shoot regeneration frequency and shoot number index revealed that all genotypes followed the same pattern for the incubation conditions tested, indicating no interaction between the factors. In both variables, a photoperiod regime responded at least 3-6 times higher than treatments with a dark pre-incubation. The degree of response depended on the genotype. No relation seems to exist between callus extension and shoot formation for the incubation conditions tested. The findings on shoot regeneration are in general agreement with other studies in which a pre-incubation in the dark did not promote regeneration. Gammoudi et al. (2017) found that a dark pre-incubation was not effective in regenerating four pepper cultivars. In *Petunia hybrida* cv. R27, it had a detrimental effect on shoot regeneration frequency (Reuveni and Evenor 2007). In lavandula (*Lavandula latifolia* Medicus), it was not beneficial on the frequency of bud and shoot regeneration when a high auxin concentration (6.0 or 11.0 µM) was used in the induction medium (Calvo and Segura 1989). Unlike, a dark pre-incubation was essential for optimal frequency of embryos or shoots in cucumber (Punja et al. 1990a). Likewise, it resulted in optimal shoot frequency and number of shoots per explant in chrysanthemum (Naing et al. 2015), and in black locus (*Robinia pseudoacacia* L.) (Arrillaga and Merkle 1993).

The mechanisms underlying the effects of a dark pre-incubation on *in vitro* morphogenesis are complex and poorly understood.

By pre-incubating in the dark, tissues could experience a redirection of resources, a change in the levels of endogenous growth regulators, or an altered sensitivity to growth regulators (Zobayed and Saxena 2003). Light and dark regimes influenced hormonal balance needed for efficient regeneration (Calvo and Segura 1989). A short light exposure on seedlings grown in the dark reduced the growth rate and altered the ratio of free to conjugated IAA (Bandurski et al. 1977). In the initiation of shoots in light-and dark-grown tobacco callus, the ethylene produced in dark culture was much higher (Huxter et al. 1981). A link has been established between ethylene, responses to stress, and the ability to regenerate (Neves et al. 2021). Light conditions also play a role in the biosynthesis of secondary metabolites (Mir et al. 2017). Different secondary compounds are known to modulate *in vitro* plant morphogenesis (Chattopadhyay 2017). For example, some phenolic compounds regulate IAA degradation, phenylpropanoids interact with auxins and act antagonistically to gibberellins, and the role of flavonoids as auxin transport inhibitors (Brown et al. 2001; Chattopadhyay 2017, and references therein). Light regimes can also influence the cell cycle (Halaban 1972; Stirk et al. 2014). Reuveni and Evenor (2007) reported a genetic component for regeneration in darkness or light in species of petunia.

Further molecular, genetic and physiological studies are needed to understand the role of light on *in vitro* morphogenesis, how it interacts with other elements, and how its effects are mediated.

## 5. Conclusions

*In vitro* shoot regeneration of one inbred line and two commercial cultivars of cucumber was significantly higher when, in shoot induction, incubation was performed under photoperiod (16L:8D h), unlike when pre-incubation in the dark followed by the same photoperiod was used. Optimized regeneration systems for selected genotypes are required to achieve a workable efficiency to apply biotechnological approaches in this species, such as large-scale micropropagation and the application of genetic transformation technologies for crop improvement, and a better understanding of its genetic basis.

## Abbreviations

BAP: 6-benzylaminopurine
IAA: Indole-3-acetic acid
KIN: Kinetin
MS: Murashige and Skoog (1962)

## Acknowledgements

The author is grateful to the Spanish Agency for International Development Cooperation (AECID) for the Ph.D. grant.

